# MSMEG_0311 is a conserved essential polar protein involved in mycobacterium cell wall metabolism

**DOI:** 10.1101/2023.09.11.557293

**Authors:** Megha Sodani, Chitra S. Misra, Savita Kulkarni, Devashish Rath

**Author notes:** Corresponding Authors: Devashish Rath, Applied Genomics Section, Bhabha Atomic Research Centre, Mumbai, India., Savita Kulkarni, Radiation Medicine Centre, Medical group, Bhabha Atomic Research Centre, Mumbai, India. Homi Bhabha National Institute, Training School Complex, Anushaktinagar, Mumbai-400094, Maharashtra, India (Megha Sodani, Savita Kulkarni, Devashish Rath).

## Abstract

Cell wall synthesis and cell division are two closely linked pathways in a bacterial cell which distinctly influence the growth and survival of a bacterium. This requires an appreciable coordination between the two processes, more so, in case of mycobacteria with an intricate multi-layered cell wall structure. In this study, we investigated a conserved gene cluster and show that knockdown of most of the genes in this cluster leads to growth defects. We further characterised one of the genes, MSMEG_0311. The repression of this gene not only imparts severe growth defect but also changes colony morphology. We demonstrate that the protein preferentially localises to the polar region and show its influence on the polar growth of the bacillus. A combination of permeability and drug susceptibility assay strongly suggests a cell wall associated function of this gene which is also corroborated by transcriptomic analysis of the knockdown. Considering the gene is highly conserved across mycobacterial species and appears to be essential for growth, it may serve as a potential drug target.

## Introduction

Mycobacteria latently infects one fourth of world population and ∼10.6 million people suffer from the active disease (Bagcchi, 2023). With the soaring cases of drug resistance in this species, mycobacterial infections have emerged as a threat across the globe (Tiberi et al., 2022). Despite the whole genome sequence being available since more than two decades, a large sub-set of difficult to assay, essential genes (∼20% of genome) of mycobacteria remain uncharacterised (A. K. Singh et al., 2016). It is imperative to probe essential gene functions not only to understand the physiology of the pathogens better but also to facilitate drug development. However, the technical challenges associated with studies of essential genes and the difficulty in handling of a slow growing infectious agent have led to their poorer characterisation (Peters et al., 2016). In this context, the recently developed programmable Clustered Regularly Interspaced Short Palindromic Repeats interference (CRISPRi) system has proved to be an effective tool to interrogate essential genes in bacteria including mycobacteria (Agarwal, 2020; Bindal, Krishnamurthi, Seshasayee, & Rath, 2017; Qi et al., 2013; A. K. Singh et al., 2016). Additionally, the availability of a surrogate model like *Mycobacterium smegmatis* helps in accelerating progress.

The unique multi-layered cell wall is a remarkable feature of the mycobacterium species, distinguishing it from other common gram positive and gram negative bacteria. This complex cell wall acts as an efficient barrier from hostile external milieu for the bacillus and it is not surprising that a variety of important drugs target different cell wall components. The understanding of many facets of cell wall synthesis in mycobacteria is far from complete. There is a plethora of unsolved puzzles, for example, trafficking of various lipids across the multi-layered cell wall and role of key players in various cell wall pathways which need to be unveiled. A deeper understanding of the complex cell wall components and their biosynthetic pathways is likely to reveal additional potential drug targets. In this work, we study a conserved mycobacterial gene cluster, consisting mostly of essential genes. The knockdown of these genes results in a severe growth defect phenotype in *M. smegmatis*. Based on the observed extreme phenotype, MSMEG_0311 was studied further. We present evidence to show that it influences cell permeability and has a role in polar cell growth. Overall, our results point towards an essential cell-wall associated role for MSMEG_0311.

## Results

### MSMEG_0311 is located in a genomic locus that is highly conserved in mycobacteria

MSMEG_0311 is an uncharacterized gene of unknown function. The domain analysis of the ORF via InterPro-Pfam (Mistry et al., 2020) was performed and hits matching glycosyltransferases were received. The N-terminal domain (28-193 amino acids) of the protein has been annotated as a glycosyltransferase family 4 protein (GT-4), which catalyzes the transfer of sugar moieties from activated donor molecules to specific acceptor molecules (Fig. 1A). The alpha fold protein structure available for this protein has the GT-B topology with two Rossmann folds connected by a short linker (Fig. 1B), one of the two protein topologies observed for nucleotide-sugar-dependent glycosyltransferases (Jumper et al., 2021). While, GT-B enzymes do not seem to share any strictly conserved residues, they are often found to contain the conserved EXE motif (Glu-X_7_-Glu) (Absmanner, Schmeiser, Kämpf, & Lehle, 2010; Rosén, Edman, Sjöström, & Wieslander, 2004). This motif has been postulated to be associated with the retention of C1 configuration in various retaining glycosyltransferases. *Mycobacterium tuberculosis* has seven such GT-4 family proteins (Berg, Kaur, Jackson, & Brennan, 2007). Their homologs in *M. smegmatis* were searched and investigated for their sequence conservation with MSMEG_0311 (Fig. 1C). The presence of this motif in other alpha glycosyltransferases further fortifies the associated glycosyltransferase activity of MSMEG_0311.

**Fig.1.**
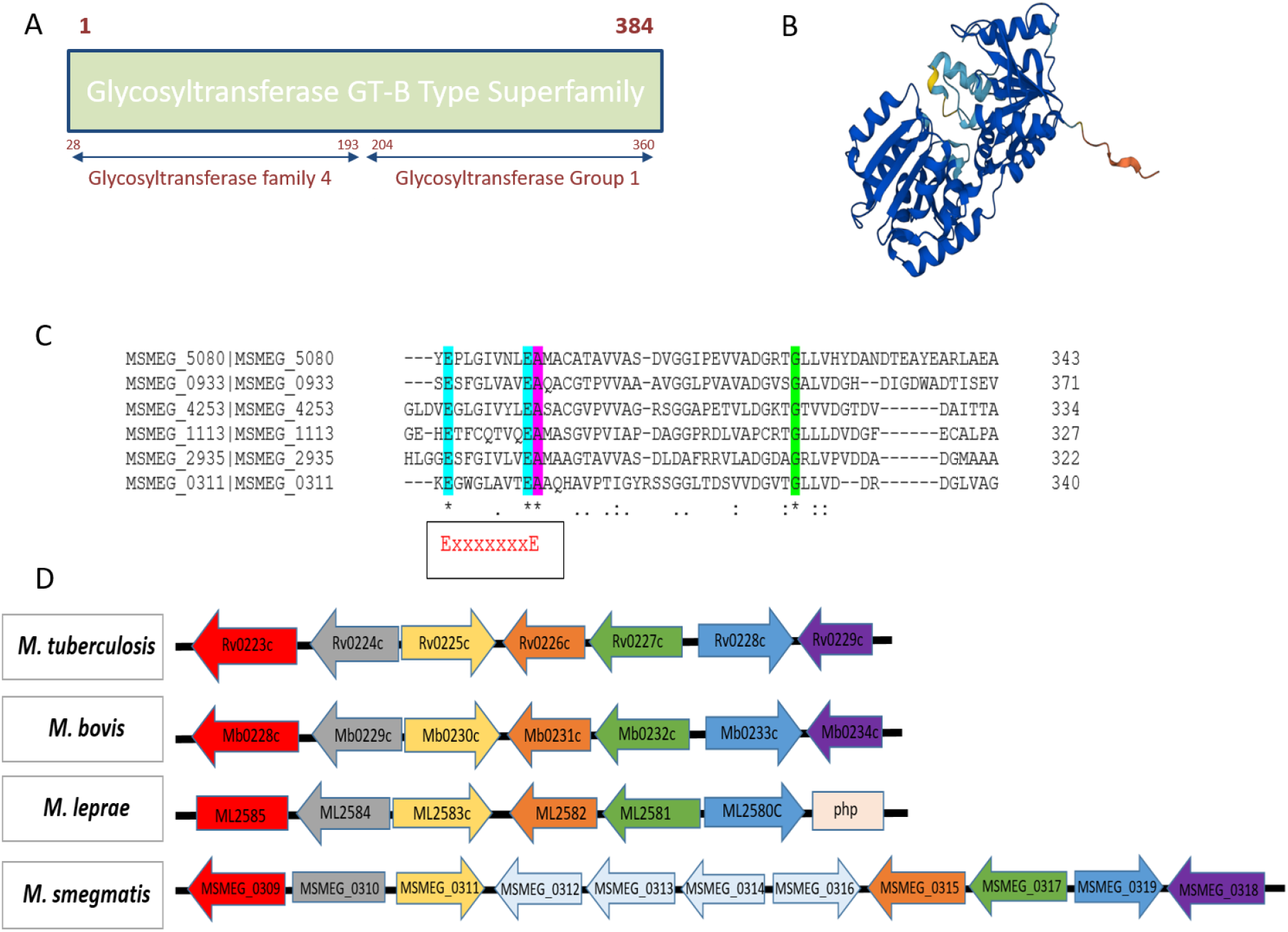
Genomic organisation and conservation of MSMEG_0311. A) Schematic of domain analysis of MSMEG_0311 showing the presence of glycosyltransferase domain. B) Alpha fold protein structure of MSMEG_0311 showing the Rossmann fold. C) Multiple sequence alignment of MSMEG_0311 with other known glycosyltransferase -GT4 showing conservation of EXE_7_ motif and other conserved residues highlighted in green. D) Genomic organisation of MSMEG_0311 and its homologues in different genus of mycobacterium (the direction of arrow represents the direction of the frame; the boxed genes are annotated as pseudogenes).

We examined the genomic locus encoding MSMEG_0311 and its orthologues across different mycobacterial species (Table S1). All its orthologues are encoded as a mono-cistron, with almost an identical organisation of flanking genes (Fig. 1D). This gene is conserved across mycobacterium species, such as *M. tuberculosis*, *M. bovis*, and *M. leprae* (Fig. 1D). It was interesting to note the conservation of the homologue even in *M. leprae* which has undergone reductive evolution and has several genes deleted from its genome and also several accumulated as pseudogenes (Gómez-Valero, Rocha, Latorre, & Silva, 2007; P. Singh & Cole, 2011). MSMEG_0311 gene homologue of *M. leprae* is not a pseudogene and is annotated as a putative glycosyltransferase (Gómez-Valero et al., 2007). The analysis of the neighbouring genes in this region showed an overall similar genomic organisation in these genomes. Except for the insertion of few genes, the gene organisation was similar in *M. smegmatis* (Fig. 1D). The synteny and conservation of MSMEG_0311 points towards an important role for this gene in mycobacteria.

### Knock-down of MSMEG_0311 and its neighbouring genes significantly restricts growth in vitro

A survey of MSMEG_0311 neighbouring genes showed that most of them are poorly characterised. In order to understand their role in growth, MSMEG_0311 and its neighbouring genes which were conserved across different mycobacterial species viz. MSMEG_0310, MSMEG_0315, MSMEG_0317 and MSMEG_0319 were targeted with CRISPRi. For knocking down expression of the genes, an *M. smegmatis* strain with *cas-12a* (Fig. S1) integrated in the genome under a tetracycline inducible promoter was constructed. The strain was transformed with plasmids expressing crRNAs (containing spacers targeting selected genes on the template strand) from a constitutive promoter. A plasmid expressing crRNA containing a non-targeting spacer NTA (Non-Targeted Array, a scrambled sequence with no complementarities to the mycobacterium genome) was used as control.

The transformation efficiency of plasmids encoding spacers targeting MSMEG_0310, MSMEG_0311, MSMEG_0317 and MSMEG_ 0319 was markedly reduced compared to the control plasmid even in the absence of the inducer, Anhydrotetracycline (ATc) (Fig. 2A). Induction of the CRISPRi with ATc further reduced the transformation efficiency of plasmids targeting MSMEG_0310 and MSMEG_0311. Transformation efficiency of the plasmid targeting MSMEG_0315 was similar to the control under both inducing and non-inducing conditions (Fig. 2A). To rule out a spacer-specific effect, three different spacers targeting different regions of the MSMEG_0315 ORF were employed, none of which showed a change in transformation efficiency (Fig. S2A) as compared to the control plasmid.

**Fig.2.**
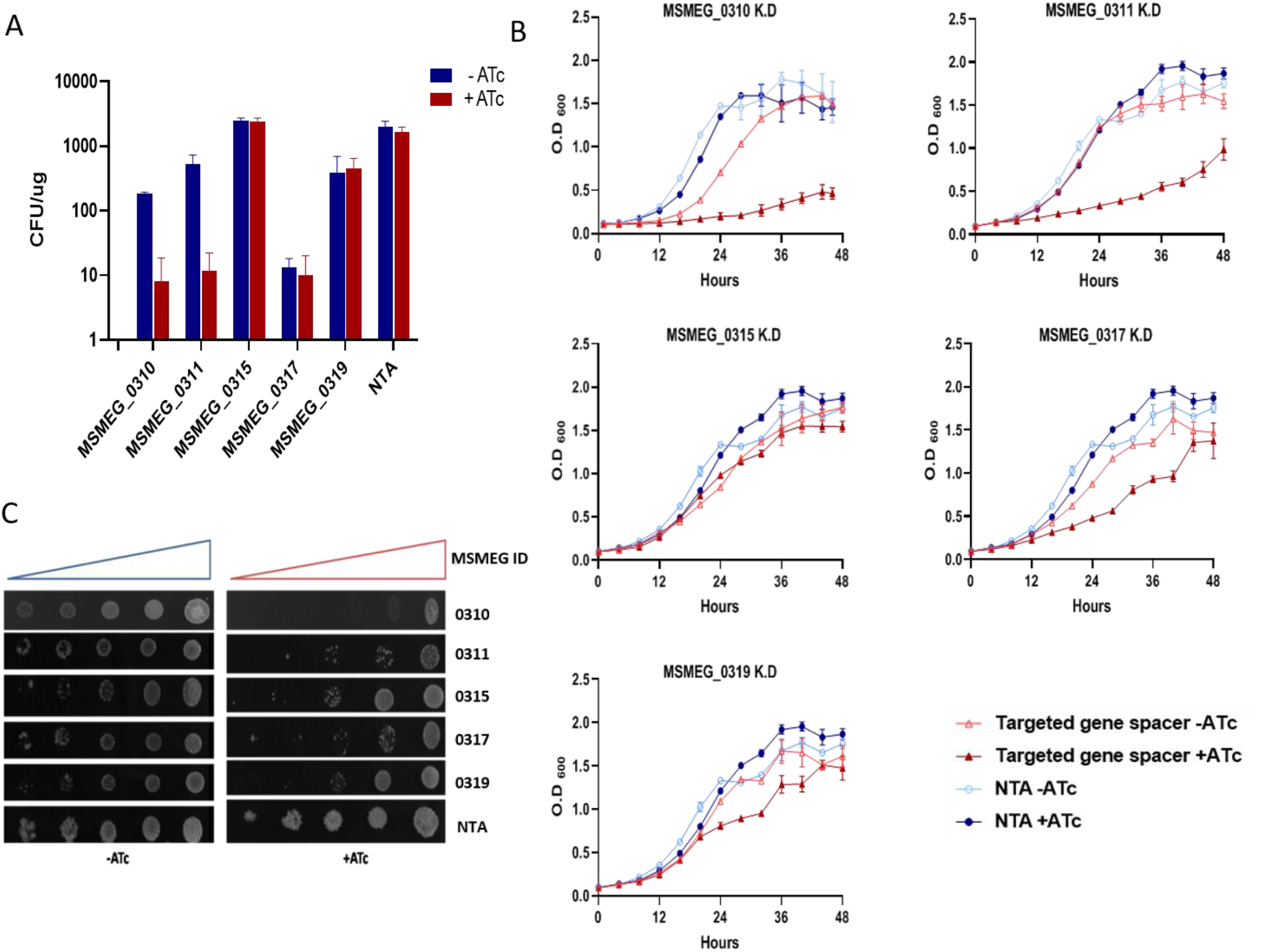
Growth phenotype of knockdown strains of MSMEG_0310, MSMEG_0311, MSMEG_0315, MSMEG_0317 and MSMEG_0319. A) Transformation efficiency of plasmids bearing spacers targeting the given genes and control NTA when introduced into *M. smegmatis* strain bearing Cas12a effector protein. B) Growth curves of the knockdown strains in broth conditions in presence and absence of inducer, ATc. C) Growth of knockdown strains spotted on solid medium in presence and absence of inducer, ATc. Equal volumes of serial dilutions were spotted.

Subsequently, we did a growth assay and evaluated the fitness of the targeted strains as compared to the non-targeted control in presence and absence of ATc in broth and agar conditions. MSMEG_0311 targeted strain showed a severe growth defect only under inducing conditions (Fig. 2B and 2C). MSMEG_0311 knock down strain (MSMEG_0311 K.D) not only displayed a longer lag phase but also a marked reduction in growth rate compared to the NTA strain (Fig. 2B) upon induction of CRISPRi. The severe reduction in growth suggests that the gene is essential in *M. smegmatis*. Additionally, an MSMEG_0311 over-expressing strain showed no growth defect and behaved like the control (Fig. S2B). Further, we targeted its homologue (*Mb0230*) in *Mycobacterium bovis* and a similar degree of growth defect was seen suggesting the essentiality of gene in a slow growing species, *M. bovis* as well (Fig. S2D). The orthologue of MSMEG_0315 in *M. tuberculosis* is annotated as an essential gene based on transposon studies (Griffin et al., 2011; Sassetti, Boyd, & Rubin, 2003). However, knockdown of MSMEG_0315 in *M. smegmatis* showed no growth defect in both agar and broth conditions (Fig. 2B and 2C). The contrasting observations could be explained by the fact that CRISPRi could have resulted in residual levels of the protein that might have been sufficient for supporting growth, while transposon insertion would have resulted in a complete knock-out of MSMEG_0315, alternatively the gene homologue may be essential in *M. tuberculosis* but not in *M. smegmatis*.

Depletion of MSMEG_0317 resulted in a moderate growth defect without induction but a severe growth defect in presence of the inducer (Fig. 2B and 2C). This result is in conformity with a recently published report (Bosch et al., 2021; Gupta & Gwin, 2022). MSMEG_0319 knockdown strain showed a growth defect on solid medium, forming tiny colonies on agar plates (Fig. S2C). However, the growth of the knock down strain was not much affected compared to the control NTA strain in broth (Fig. 2B). Interestingly, knock-down of MSMEG_0310, which is annotated as a pseudogene (Fleischmann et al., 2006) resulted in a growth defect in both solid and liquid medium (Fig. 2B and 2C) suggesting that it produces a true functional protein. Among these genes, MSMEG_0311 which showed a severe growth defect upon knockdown was further investigated.

### MSMEG_0311 knock down strain display altered morphology

As the conserved homologues of some of the genes in the neighbourhood of MSMEG_0311 were shown to be membrane associated or involved in cell wall component synthesis in the related species, we hypothesized a cell wall associated function for MSMEG_0311 (Cashmore et al., 2017; Rainczuk et al., 2020; Yamaryo-Botte et al., 2015). The cell wall is the primary interface with the external environment, and alterations in the cell wall composition can lead to pronounced effects on colony morphology (Chen, Teng, Wen, Zhang, & Huang, 2020). In presence of the CRISPRi inducer, ATc, MSMEG_0311 K.D strain showed altered colony morphology (Fig. 3A). The colonies were smooth and lacked the discrete cording which is the hall mark of mycobacterium colony morphology. The control strain bearing the NTA spacer displayed colony morphology similar to the wild type *M. smegmatis* with serrated colony boundary and a rough surface with distinct cording (Fig. 3A). The cording feature of the colony is attributed to presence of trehalose dimycolate (TDM) (Hunter, Venkataprasad, & Olsen, 2006) and it is interesting to note that MSMEG_0317, a neighbouring gene, has been recently reported to play role in transport of trehalose conjugated mycolic acid across the inner membrane (Gupta & Gwin, 2022).

**Fig.3.**
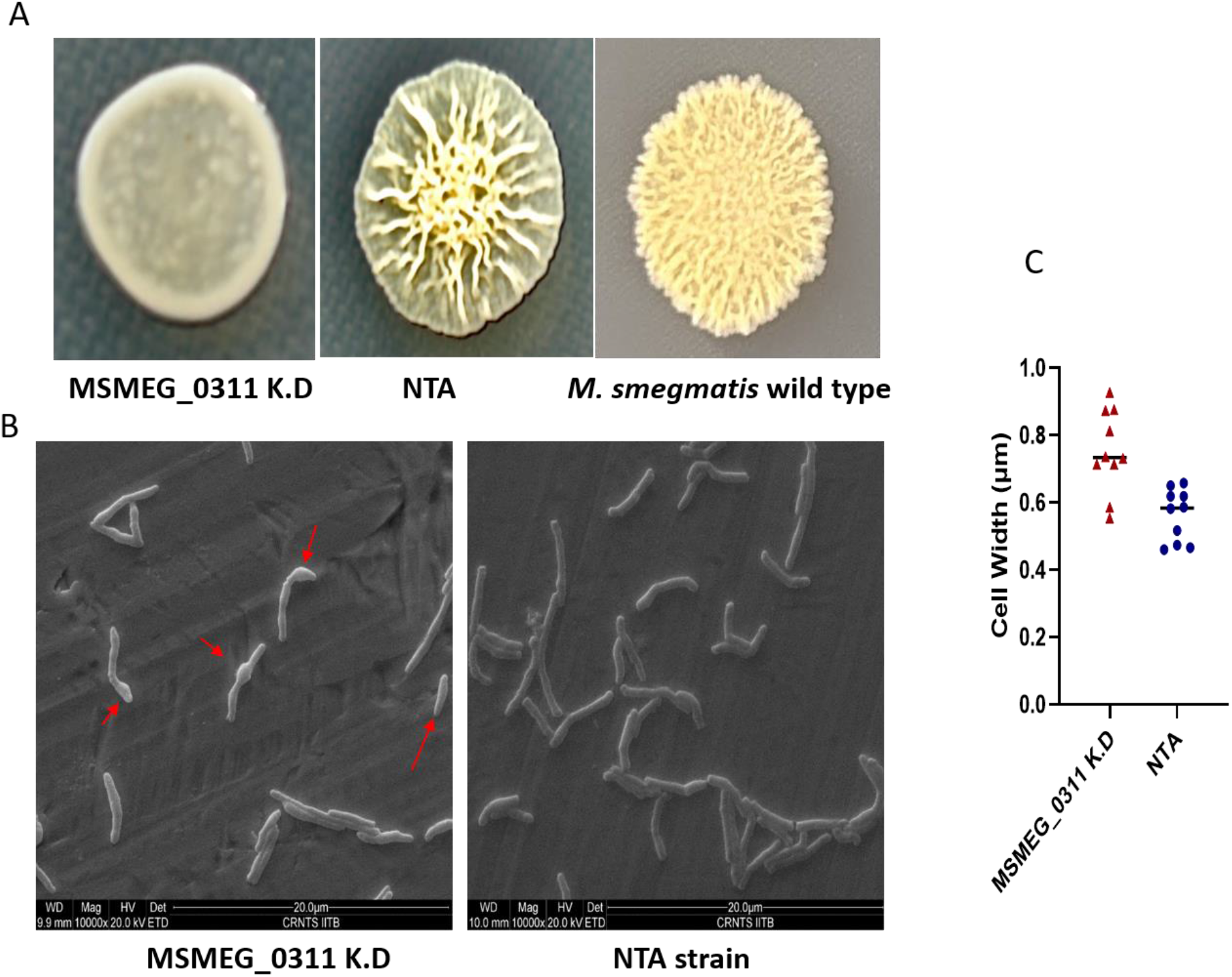
Morphological defects seen in MSMEG_0311 K.D strain. A) Colony morphology of MSMEG_0311 K.D strain, NTA and wild type *M. smegmatis.* B) Electron micrographs of MSMEG_0311 K.D strain and control NTA showing deformed crooked cells in the former (red arrows). C) Quantitative analysis of the effect of knockdown on cell width using ImageJ.

The MSMEG_0311 K.D strain was viewed under scanning electron microscope. It was observed that the surface morphology was altered with many cells displaying bulging at peripolar and mid septum region (Fig. 3B and 3C). Though only a fraction of the cells showed the phenotype, such population was totally absent in the control strain. As cell wall synthesis and cell division are often intertwined, the bulging of MSMEG_0311 depleted cells points towards a cell wall synthesis or cell division related role of MSMEG_0311.

### Knock-down of MSMEG_0311 affects membrane permeability

MSMEG_0311 K.D strain was tested for membrane permeability by SDS disc diffusion assay and susceptibility to the surfactant was scored in terms of the zone of inhibition (ZOI). SDS is a well-known anionic surfactant, routinely used to test mycobacterium membrane integrity (Bharti et al., 2021). Upon exposure to 2.5% SDS, MSMEG_0311 K.D strain showed a significantly larger ZOI compared to control, NTA strain (Fig. 4A). While MSMEG_0311 K.D exhibited a ZOI of 33 ± 2 mm, the NTA ZOI was measured at 26 ± 1 mm (average of three independent readings) (Fig. 4B). As observed, MSMEG_0311 K.D strain was significantly more susceptible to cell envelope stress caused by SDS than the NTA strain.

**Fig.4.**
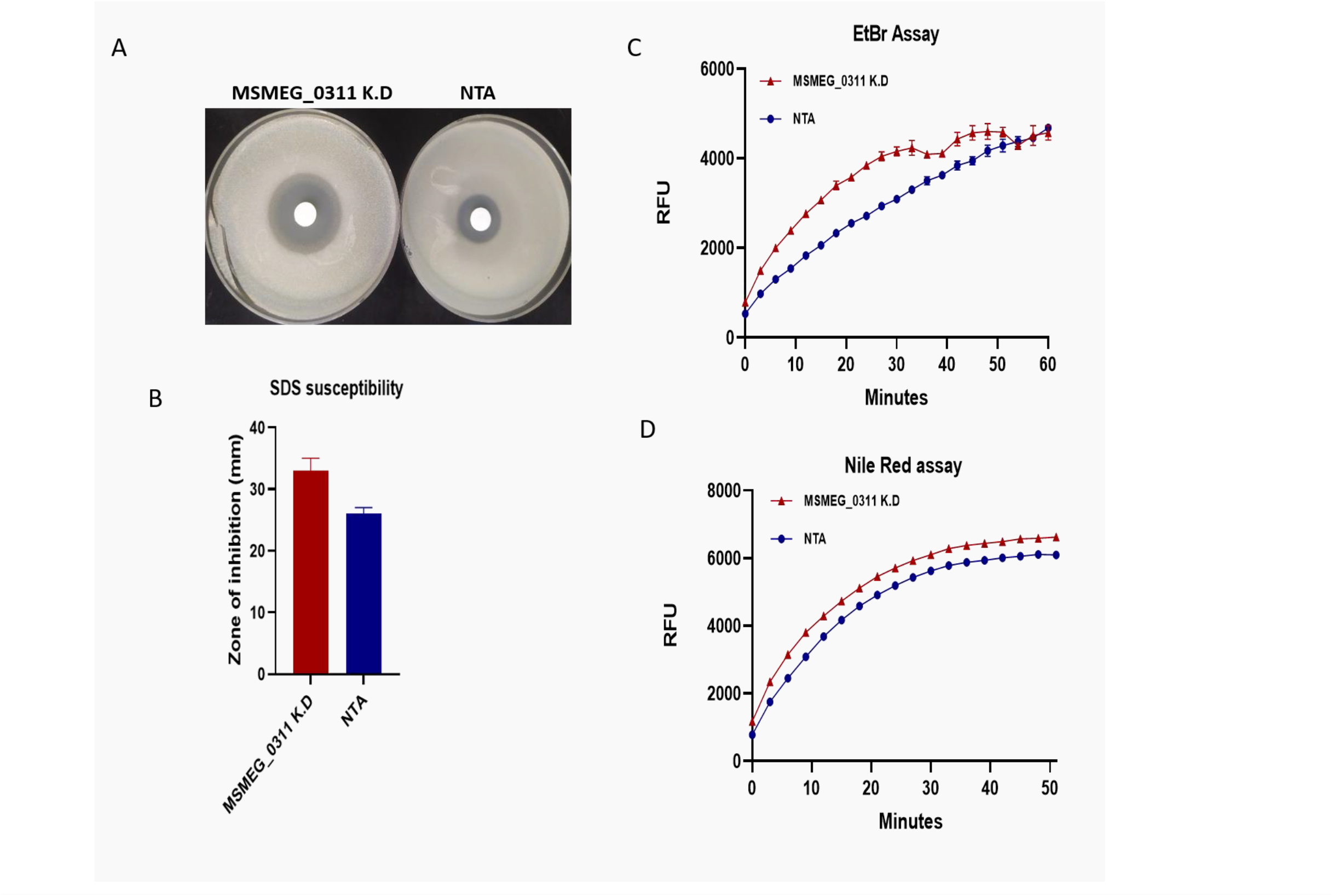
Effect on cell permeability of MSMEG_0311 K.D strain. A) SDS disc assay showing zone of inhibition of MSMEG_0311 K.D and NTA strain. B) Quantitative analysis of ZOI of MSMEG_0311 K.D and NTA strain. C) EtBr influx assay showing higher uptake of dye in MSMEG_0311 K.D strain. D) Nile red influx assay showing higher uptake of dye in MSMEG_0311 K.D strain.

To evaluate cell wall permeability, ability to take up the dyes ethidium bromide (EtBr) and nile red was assessed (Abdalla et al., 2020). EtBr is weakly fluorescent when free in the medium, but becomes fluorescent when intercalated in DNA. Similarly, nile red fluoresces upon interacting with intra-cellular lipids. MSMEG_0311 K.D and the NTA strains were subjected to EtBr and nile red treatment and the fluorescence kinetics was monitored using plate reader. MSMEG_0311 K.D strain accumulated both EtBr and nile red in higher amounts compared to control NTA (Fig. 4C and 4D) suggesting increased cell permeability in the knockdown strain. Though there was a moderate increase in uptake of the dye, it was significant as confirmed with three independent replicates. In summary, depletion of MSMEG_0311 in *M. smegmatis* significantly altered its cell wall associated properties such as colony morphology, permeability and surfactant sensitivity.

### Transcriptome analysis of MSMEG_0311 K.D strain

The genome wide effects of MSMEG_0311 knock-down was analysed with RNASeq. A total of 995 genes were differentially expressed, with 596 genes being up-regulated and 399 found to be down-regulated (P-Value ≤ 0.05 and Log2Fc ≥1 and ≤-1, data not shown) in comparison to the control NTA. Most of the differentially expressed genes belonged to the categories of conserved hypothetical proteins, cell wall processes, and intermediary metabolism. The genes, which showed maximum change in expression were mostly uncharacterised genes of unknown functions like-MSMEG_6606, MSMEG_6608, MSMEG_6579 (Fig. 5A). Among the top 20 differentially expressed genes, those belonging to ABC cassette family such as MSMEG_0113 (*tauC*-sulphur metabolism), MSMEG_0116 (*cysA1*-sulphate transport) and MSMEG_2015-2016 (*modB*-molybdenum transporter) were down-regulated, while MSMEG_6766 (antibiotic transporter ATP-binding protein) was upregulated (Fig. 5A).

**Fig.5.**
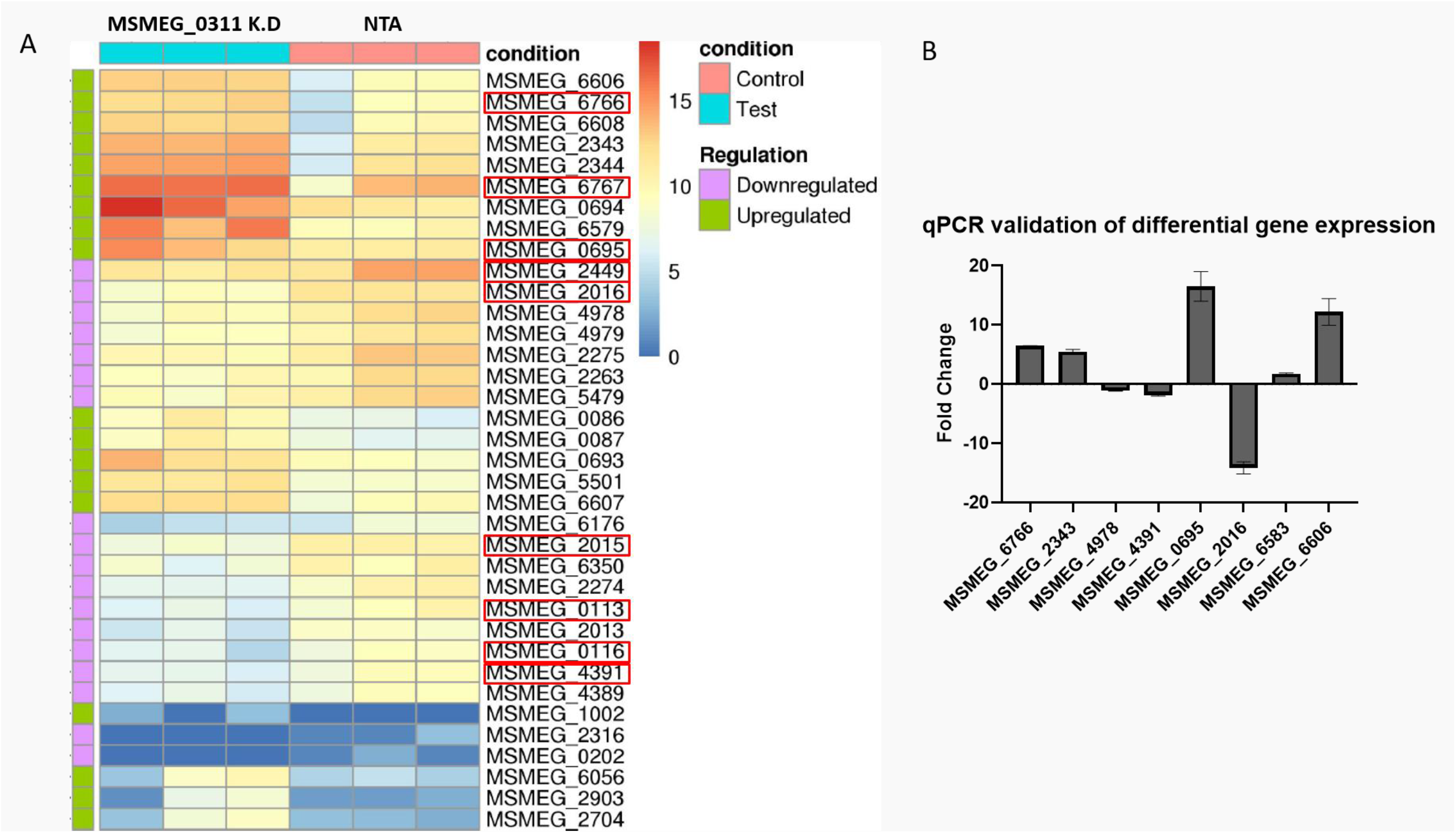
Transcriptome analysis of MSMEG_0311 K.D strain. A) Heat map depicting expression of the top most differentially expressed genes in MSMEG_0311 K.D strain vs NTA listed as per statistical significance (genes are known to be involved in cell wall or lipid metabolism are highlighted). B) Quantitative analysis of RNA levels by qRT-PCR of some differentially expressed genes detected in RNASeq.

Interestingly, the RNAseq data also showed upregulation of *iniA* gene (MSMEG_0695) which was among the top 20 gene set (Fig. 5A). The expression of seven of the genes shown to be differentially expressed by RNA-Seq analysis was validated using qPCR (Fig. 5B). Among these, *iniA* was found to be highly up-regulated (∼17-fold) in the MSMEG_0311K.D strain. *iniA* and *iniB* are part of *iniBAC* operon which is reported to be up-regulated in in response to isoniazid or ethambutol exposure in *M. tuberculosis* (Colangeli et al., 2005). On closer observation, we also noticed that several SigF regulated genes (Hümpel, Gebhard, Cook, & Berney, 2010) like *cysA2,*MSMEG_2343, MSMEG_ 2344, MSMEG_6579 and MSMEG_6767 were upregulated. SigF is an alternative sigma factor that is highly conserved among species of the genus-Mycobacterium and is known to modulate the cell surface architecture and lipid biosynthesis (A. K. Singh et al., 2015). Additionally, it was observed that several cell wall associated genes e.g., MSMEG_2343, a methylesterase that is part of a two-component Rieske oxygenase (RO) in the cholesterol catabolic pathway of Mycobacterium, MSMEG_6767 coding for mycocerosic acid synthase, MSMEG_2449 coding formethyl malonate-semialdehyde dehydrogenase, MSMEG_2013 (*fabG3*) and MSMEG_4391 (*fadE13*) known to be involved in lipid synthesis, and pks15 showed altered expression in the knockdown strain. Collectively, the transcriptome data strongly suggests the involvement of MSMEG_0311 in cell wall metabolism. The pathway association of this gene needs to be further investigated at depth to determine its specific function.

### MSMEG_0311 is involved in cell wall associated function

Genes involved in cell wall metabolism characteristically influence antibiotic tolerance (Boutte et al., 2016; Geisinger et al., 2020; Shamma et al., 2021). We sought to examine the antibiotic susceptibility of the MSMEG_0311 K.D strain. As MSMEG _0311 appeared to be an essential gene, the susceptibility assays were designed such that the viability loss due to MSMEG _0311 per se was already accounted for. To test the effect of the knock-down on permeability of the cell wall, the strains were cultured as lawn on agar medium and exposed to paper discs impregnated with a variety of antibiotics. The sensitivity of the strains was assessed by observing the size of zone of inhibition (ZOI) around the discs. We tested several antibiotics targeting different cellular pathways like protein synthesis, cell wall metabolism and energy and respiration. We observed that MSMEG_0311 K.D strain was uniformly more sensitive to all the protein synthesis inhibitors (Fig. 6A and S3). The results were variable with respect to antibiotics targeting the cell wall. While it was slightly less susceptible to isoniazid compared to the control strain, NTA, it did not show any differential sensitivity towards beta-lactam class antibiotics imipenem and ampicillin (data not shown). Nevertheless, in case of ethambutol that targets arabinogalactan synthesis, MSMEG_0311 K.D showed a lower MIC in agar disc assay (Fig. 6A and S3).

**Fig.6.**
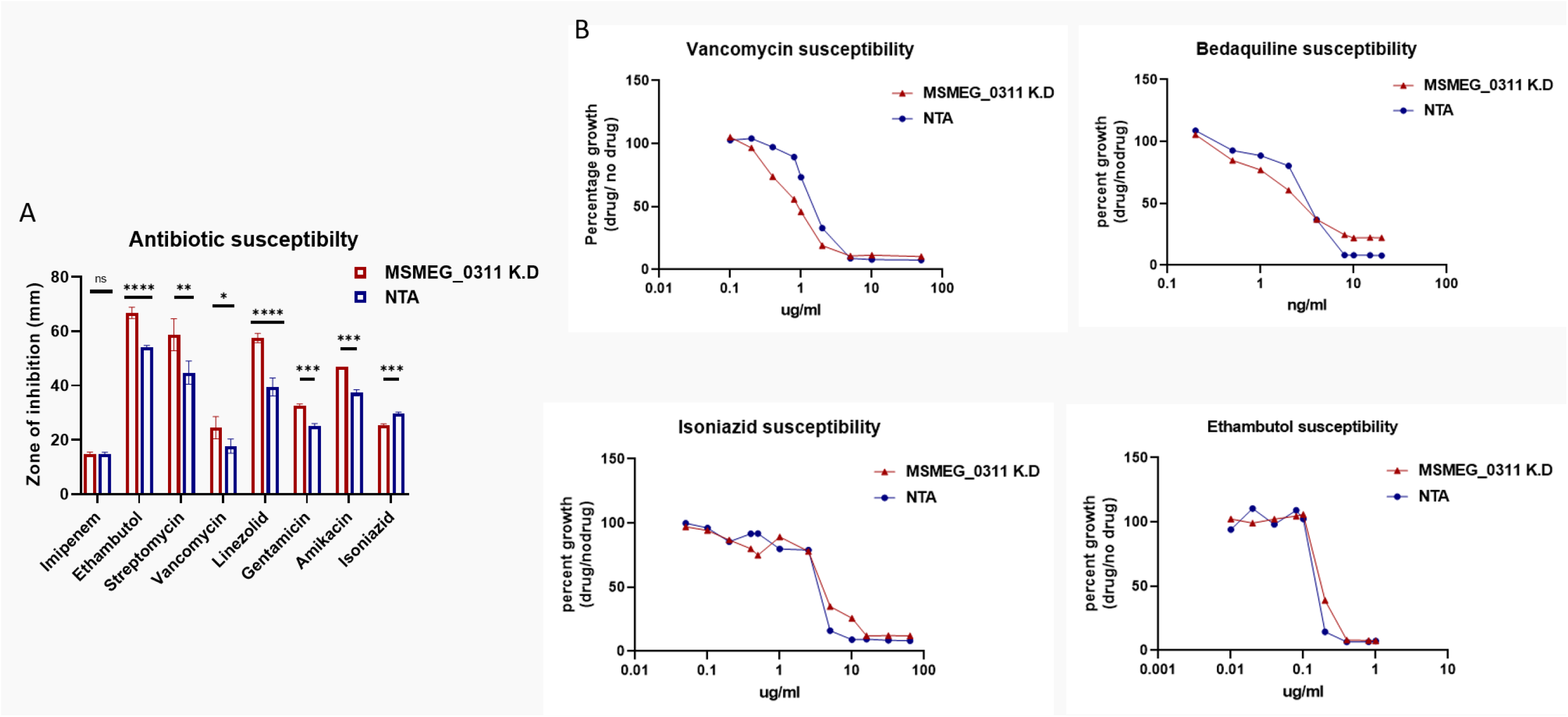
Antibiotic susceptibility of MSMEG_0311 K.D strain. A) Antibiotic disc assay of MSMEG_0311 K.D strain depicting sensitivities (ZOI) against given antibiotics. B) Antibiotic broth assay showing the susceptibility of MSMEG_0311 K.D strain against each drug at varied concentrations. The effect on growth due to repression of gene itself was normalised by plotting the percent growth in media containing the drug compared to growth in media without any drug at various concentration of antibiotics.

A broth assay was also developed to investigate the differential susceptibility as genes playing a role in cell wall metabolism behave differently in agar and broth conditions. The drug susceptibility was examined at different concentrations spanning the predicted minimum inhibitory concentration and the growth reduction was normalised to control strain without the exposure to the given drug. Collectively, we observed no differential sensitivity towards isoniazid and ethambutol in broth assay, while higher sensitivity was observed to vancomycin, and to bedaquiline at lower concentrations (Fig. 6B). Overall, we saw differential level of antibiotic sensitivities in agar and broth conditions.

### MSMEG_0311 localises to the poles of growing cells

Analysis of MSMEG_0311 by TBpred predicted it to be a cytoplasmic protein with no signal peptide (Rashid, Saha, & Raghava, 2007). To determine the sub-cellular location of MSMEG_0311, it was tagged at the C-terminal with a fluorescent protein and constitutively expressed under the control of a heat shock protein promoter (Phsp). To reduce steric interference between the green fluorescent protein (GFP) and the C-terminal of MSMEG_0311, a 10-amino acid linker (ASGSAGSAGSA) was added between the two. Subsequently, a FLAG tag was added at C-terminus of the fusion protein. *M. smegmatis* was transformed with the vector coding for the chimeric protein. Expression was confirmed with an anti-FLAG antibody in western blot where the fusion protein showed an approximate size of ∼71 kDa which is in agreement with the expected size (MSMEG_0311 ∼ 43 kDa, GFP ∼27 kDa and FLAG tag ∼ 1 kDa) (Fig. 7A). Fluorescence micrographs showed that MSMEG_0311-GFP localizes to the poles of mycobacterial cell (Fig. 7B). Interestingly, one pole was found to be brighter than the other indicating the differential accumulation of MSMEG_0311 to the old or the new pole (Fig. 7C). Control experiments revealed that untransformed *M. smegmatis* mc^2^155 displayed no fluorescence (data not shown), whereas expression of GFP alone resulted in a distributed fluorescence throughout the cells (Fig. 7B).

**Fig.7.**
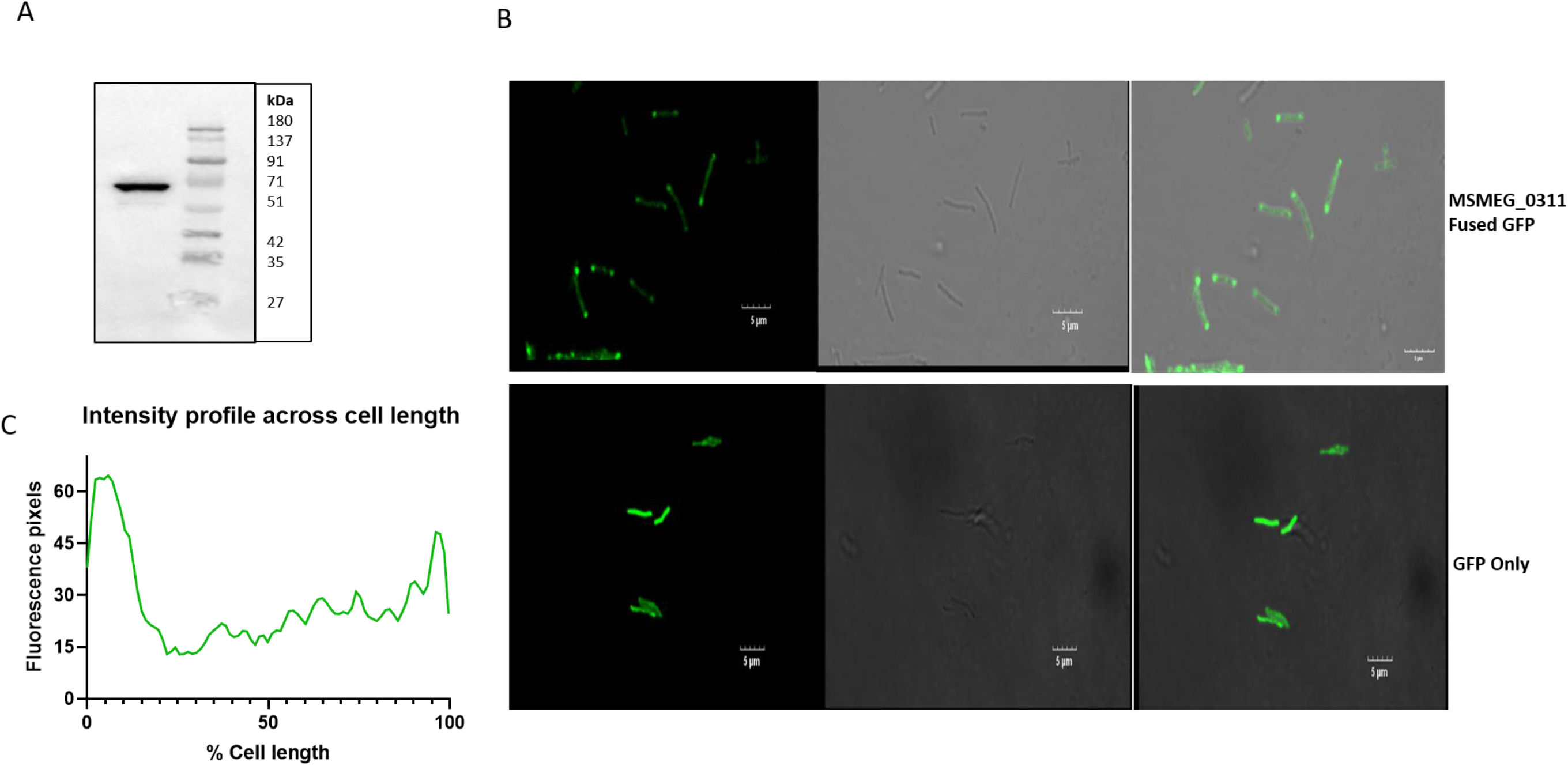
Polar localisation of GFP protein. A) Western blot showing the expression of GFP fused to MSMEG_0311 protein. B) Fluorescence micrographs showing accumulation of MSMEG_0311 at the poles (left panel: GFP filter, middle panel: Differential Interference Contrast (DIC), right panel: Merged). C) Intensity profile showing differential levels of protein at the two ends of the cell.

Emerging evidence indicates that mycobacteria organize their cellular structures differently than in established models (Aldridge et al., 2012; Rego, Audette, & Rubin, 2017). Mycobacteria incorporate nascent cell wall materials exclusively at the cell poles and septa (Meniche et al., 2014). As mycobacterium divides asymmetrically and the site of cell wall synthesis is confined towards poles rather than along the lateral axis, localization of MSMEG_0311 at poles strongly hints towards a cell wall associated function for this protein.

### MSMEG_0311 influences polar growth in mycobacteria

To visualise the effect of MSMEG_0311 on cellular growth polarity, we employed fluorescent D amino acid labels (RADA dye) which label nascent peptidoglycan (PG) and are therefore an ideal tool to study spatial aspects of PG synthesis (Kuru, Tekkam, Hall, Brun, & Van Nieuwenhze, 2015). The dye gave us an opportunity to monitor bacterial cell wall synthesis in mycobacteria which grow asymmetrically from poles rather than along the swath (García-Heredia et al., 2018). The dye was used as a probe to visualise the fraction of new growth added in a cell to demarcate old pole from new (Fig. 8A). In a pulse chase experiment, the strains were exposed to the RADA dye long enough for saturated uptake in whole cell and the dye was removed during the chase period. We probed the GFP fused MSMEG_0311 over-expressing strain with the RADA pulse and found that the cell continues to divide asymmetrically with one pole growing more than the other (Fig. 8B). Interestingly, we discovered that the brighter pole was specifically localised to the old pole where the protein likely accumulates to build up new cell wall (Fig. 8B). Being specifically localised to the growing old pole, we further investigated the role of this protein in asymmetric growth. Hence, we subjected the MSMEG_0311 K.D strain to the RADA pulse. A knockdown strain of *mmpl* served as a positive control as it is known to influence polar asymmetric growth in mycobacteria (Gupta & Gwin, 2022). It was observed that in absence of ATc, MSMEG_0311 K.D and *mmpl* K.D strain grew asymmetrically from one pole, as expected (Fig. 8C). In presence of ATc, with the depletion of respective proteins, the *mmpl* K.D strain grew almost equally from both the poles but MSMEG_0311 K.D continued to grow asymmetrically (Fig. 8C). This shows that unlike *mmpl*, the MSMEG_0311 though being a polar protein preferentially localising to the growing pole, doesn’t influence asymmetric cell growth in *M. smegmatis*.

**Fig.8.**
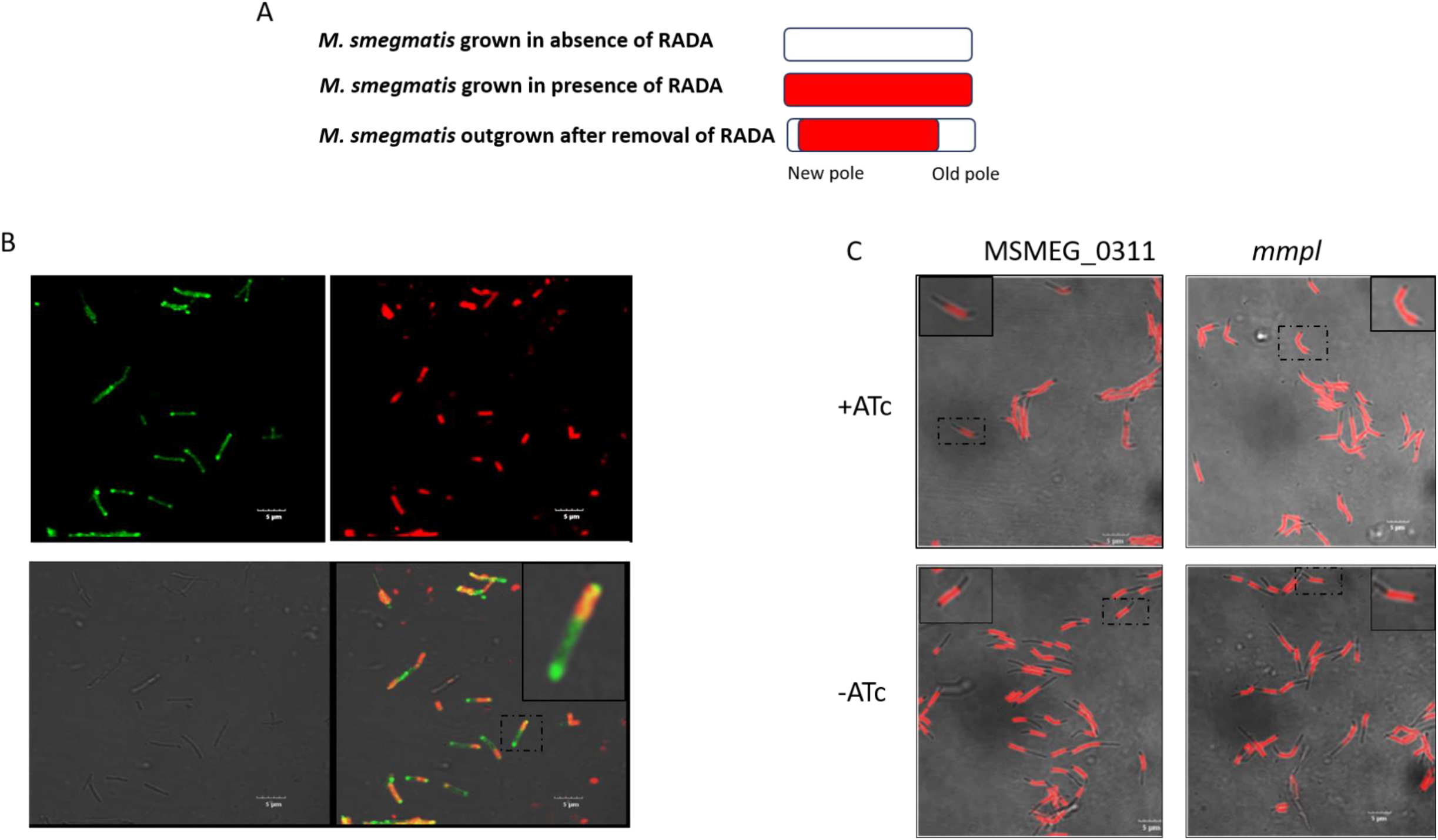
Role of MSMEG_0311 in polar growth. A) Schematic representation of asymmetrical growth of *M. smegmatis* B) Pulse chase micrographs showing localisation of MSMEG_0311 in over-expressing strain on the growing new pole (Upper left panel: GFP filter, upper right panel: Tritc filter, lower left panel: DIC, lower right panel: Merged) (the dotted box area of micrograph is zoomed in and displayed as full-lined boxes in corner) C) Pulse chase micrographs of MSMEG_0311 K.D and *mmpl* K.D strain tracking asymmetrical growth (Merged of DIC and TRITC filter) (the dotted box area of micrograph is zoomed in and displayed as full-lined boxes in corner).

## Discussion

The mycobacterium cell envelope can be partitioned into three distinct components. The outermost capsular layer is highly rich in glycans-comprising mannans, alpha-glucans, arabinomannans which are not usually found in other gram positive/negative bacteria. Beneath the capsular layer is the cell wall which has a tripartite mAGP arrangement encompassing the mycolic acid layer, arabinogalactan layer and the peptidoglycan layer. Further down, spanning the periplasmic space is the conventional plasma membrane. With such diverse components and pathways, mycobacterial cell wall metabolism needs to be studied at depth to delineate the components and biosynthetic pathways. It is imperative to investigate more and more uncharacterised genes involved in cell wall functions which can be probed for their potential as drug targets.

Here, we use CRISPRi to investigate a locus containing a cluster of poorly characterized genes, presumably involved in cell wall synthesis, in mycobacteria. The locus comprises of MSMEG_0310, MSMEG_0311, MSMEG_0315, MSMEG_0317 and MSMEG_0319. Though, the exact function of these genes are unknown, there are reports of MSMEG_0319 and MSMEG_0317 impacting the transport of trehalose monomycolate (TMM) (Belardinelli et al., 2016; Gupta & Gwin, 2022). To the best of our knowledge, there is no literature available on the role of MSMEG_0315 and MSMEG_0311, while MSMEG_0310 is annotated as a pseudogene in *M. smegmatis.* Except for MSMEG_0315, knock-down of these genes resulted in significant growth defect. This suggested that most of the genes are essential and play an important role in normal growth. Interestingly, downregulation of MSMEG_0310, which is annotated as a pseudogene, severely affected growth suggesting that it is not only a functional gene but also has an essential role in *M. smegmatis*. Among the genes, knockdown of MSMEG_0311 caused the maximum growth defect and hence, it was investigated further. With growing appreciation of vulnerable genes among essential ones (Bosch et al., 2021), we believe that MSMEG_0311 is a highly vulnerable gene as evidenced by a fold knock down of ∼ 2 leading to severe loss of fitness (Fig. S4). MSMEG_0311 knockdown cells displayed the hallmarks of loss of cell wall associated function such as altered cell and colony morphology and decreased cell wall permeability. In addition, MSMEG_0311 was found to localize to cell poles, in accordance with accumulation of new cell wall material at the poles in mycobacterium instead of the lateral side, further validating involvement of MSMEG_0311 in cell wall synthesis. Also, MSMEG_0311 protein displayed preferential accumulation at the growing old pole without affecting asymmetry in cell division during growth.

The in-silico analysis of the protein showed presence of glycosyltransferase domains of GT4 family. The complex glycan-rich cell wall of Mycobacterium necessitates the presence of a large variety of glycosyltranferase genes in its genome. The Carbohydrate Active Enzymes database or CAZy database currently enlists 43 glycosyltransferases in *M. tuberculosis*, of which 7 are of GT4 family, (Drula et al., 2022). Similarly, *M. leprae* and *M. smegmatis* have 28 and 51 glycosyltransferases, of which 5 and 13 are of GT4 family, respectively. GT4 glycosyltransferases in *M. tuberculosis* are shown to be involved in three cellular pathways, namely– phosphatidyl-inositol mannoside(PIM)/ lipoarabinomanan (LAM) synthesis (Rv2610, Rv2188, Rv0557c) (Mishra, Batt, Krumbach, Eggeling, & Besra, 2009), glycogen-glucan/polymethylated polysaccharides (PMPS) synthesis (Rv3032, Rv1212) and in mycothiol synthesis (Rv0486) (Buchmeier & Fahey, 2006). There are no reports on the role of the seventh glycosyltransferase Rv0225c, the homologue of MSMEG_0311. In LAM biosynthesis, it is predicted that the synthesis of PIM_1/2/3/4_ occurs on the cytoplasmic side after which it is flipped through the periplasmic space where polyprenoid dependent GT-C mannosyl-transferases add mannose residues to form higher PIMs and LM/LAM. Till date, these mannosyl-transferases catalysing the formation of PIM_3/4_ are unknown with an ambiguous report of PimC identified in *M. tuberculosis* CDC1551 strain (Kremer et al., 2002). It is possible that MSMEG_0311, being a GT-B glycosyltransferase might be involved in synthesis of PIM3/PIM4. The homologue of the neighbouring gene MSMEG_0317 in *Corynebacterium glutamicum*, Ncgl2759 has also been reported to be involved in the synthesis of mature LAM (Cashmore et al., 2017). Alternatively, given the tertiary structure of MSMEG_0311 being highly similar to Rv3032, a role for MSMEG_0311 in glucan/MGLP (Methylated glucose lipopolysaccharide) synthesis may be speculated, specifically when the gene homologue of Rv3032 (MSMEG_2348) in *M. smegmatis* is a pseudogene.

With the advent of CRISPRi in mycobacteria, several remarkable high throughput studies of essential genes have been performed (Bosch et al., 2021; de Wet & Winkler, 2020; Li et al., 2022). Genetic screens and clustering analysis have demonstrated common antibiotic sensitivity across genes of similar pathway (Li et al., 2022). For example, genes playing role in LAM pathway are highly sensitive to vancomycin (Fukuda et al., 2013), ethambutol and to some degree to bedaquiline at higher concentration but not sensitive to isoniazid (Li et al., 2022). Though our drug susceptibility data demonstrate a susceptibility profile partially matching with LAM synthesis pathway, involvement of MSMEG_0311 in synthesis of PMPS or mycothiol cannot be ruled out. Overall in this study we characterize the MSMEG_0311 to be an essential gene and present several lines of evidences including microscopy, cell permeability, drug susceptibility profiling and transcriptome data to show that it has a cell wall associated function. Future studies with a lipidomics approach may be required to pin-point its role in PIM/LAM, PMPS or mycothiol biosynthesis.

## Materials and Methods

### Bacterial strains and growth conditions

*E. coli* DH5α and JM109 strains were used for vector construction. Cells were grown in Luria-Bertani broth and supplemented with appropriate antibiotics (kanamycin at 50 µg/ml, hygromycin at 150 µg/ml), wherever required. To prepare solid medium, 1.5% agar was included in the broth.

Mycobacterium strains used are derivatives of *Mycobacterium smegmatis* mc^2^155, (kind gift from Dr. Amit Singh, IISc, Bangalore). *M. smegmatis* mc^2^155 was grown in Middlebrook 7H9 broth supplemented with Tween 80 (0.05%), glycerol (0.25%) or Middlebrook 7H10/11 agar supplemented with glycerol (0.5%). Appropriate antibiotics (kanamycin 20 µg/ml, hygromycin 50 µg/ml) and the inducer, anhydrotetracycline (ATc) (100 ng/ml) were added wherever needed. All cultures were grown at 37°C with shaking at 150 rpm.

### Recombinant strain construction

All the cloning and initial vector constructions were done in *E. coli* following standard recombinant DNA procedures (Sambrook, 1998) before being introduced into Mycobacterium.

A two-plasmid system CRISPRi system was constructed to create knockdown of genes in Mycobacterium. Cas12a from *Francisella novicida* carrying the relevant D917A substitution, to construct the dead Cas12a variant, was codon optimised for *C. glutamicum*. The gene was synthesised along with the *tetO* promoter and cloned in pSTKiT (Parikh et al., 2013) using KpnI and BamHI sites. For cloning, integrative vector pSTKiT was digested at KpnI and BamHI site, gel purified and ligated to the similarly digested *cas12a* fragment. The recombinant vector (pSTKiT-Cas12) was constructed in *E. coli* DH5α and confirmed by Sanger sequencing. The vector, pSTKiT-Cas12 was transformed into *M. smegmatis*.

Construction of the vector for the expression of the sgRNA targeting the gene of interest is described below. The antibiotic resistance cassette of a replicative vector, pSTKT (Parikh et al., 2013) was replaced by a synthesised hygromycin resistance cassette by cloning it at KpnI and SpeI sites to construct pSTHT. The sequence coding for the sgRNA was cloned in replicative vector pSTHT under the constitutive promoter P_smyc_ with the provision for Golden Gate spacer cloning at BsaI site. The oligo spacers, (20bp) were designed targeting the template strand in the ORF region with flanks 5’-TAGA-3’ on the top strand and 5’-A GAC-3’ on the bottom strand. The synthesised strands (2μl of 100μM) were annealed in a 20μl reaction containing TRIS buffer pH-8 (10 mM Tris HCL, 50 mM NaCl and 1 mM EDTA) by gradual cooling from 94 ^◦^C for 4 min,75 ^◦^C for 5 min, 65 ^◦^C for 15 min to 25 ^◦^C for 20 min. The annealed oligos were diluted 10 times and ligated into pSTHT using BsaI golden gate assembly. The oligos used are listed in Table S3. The cloned spacers (pSTHT-0310 K.D, pSTHT-0311 K.D, pSTHT-0315 K.D, pSTHT-0317 K.D, pSTHT-0319 K.D, pSTHT-0250 K.D) were sequenced and subsequently transformed into electro-competent *M. smegmatis* cells bearing pSTKiT–Cas12. Briefly, *M. smegmatis* culture was expanded to an O.D_600nm_ = 0.8–1.0 (100 ml culture) and pelleted (5000*g* for 5 min). The cell pellet was sequentially washed three times with 25 ml, 12 ml and 6 ml of sterile 10% glycerol respectively. The washed bacilli were then re-suspended in 10% glycerol to a final volume of 1% of the original culture volume. For each transformation, 300 ng plasmid DNA was added to 100 μl electro-competent mycobacteria and transferred to a 2 mm electroporation cuvette. Electroporation was performed using the Gene Pulser X cell electroporation system (Biorad) set at 2500 V, 1000 Ω and 25 μF. The cells were allowed to recover in media for 3 hours. After the recovery, cells were plated on 7H11 agar supplemented with the appropriate antibiotic to select for transformants.

pMV261 (Stover et al., 1991) was used for cloning the fusion protein of MSMEG_0311 and GFP. MSMEG_0311 was amplified from *M. smegmatis* genomic DNA with primers bearing the restriction site (BamHI and EcoRI) (Table S3). The gene was cloned at the given site in pMV261 to generate plasmid pMV261-0311. Next, primers bearing the linker region as flanks were used to amplify GFP gene from pMN437 (Song, Sandie, Wang, Andrade-Navarro, & Niederweis, 2008) which was ligated into (EcoRI and HindIII site) of pMV261-0311 to obtain pMV261-0311-GFP. The construct was confirmed by Sanger sequencing and transformed into *M. smegmatis*.

### RNA isolation, qPCR and RNAseq Assay

Mycobacterium strains (3-5 O.D_600nm_) were harvested at 2000*g* for 10 min. The pellet was re-suspended in 1ml TRIzol and transferred to screw capped tubes containing lysing matrix for homogenisation. The tubes containing the cell lysate were pulsed in MP Biomed (bead beater-FastPrep-25 5G) at 6 m/sec for 45 seconds for 2 cycles with a 2 min ice-cooling step in between. The lysate with matrix beads were centrifuged at 8000*g* and the supernatant was transferred to a fresh tube. RNA was isolated from the lysate using Zymo Research RNA zol kit according to the manufacturer’s instructions. The eluted RNA was treated with TURBO™ DNase for 1 hour at 37°C to remove residual DNA and was column purified using NEB Monarch DNA clean up kit. The purified RNA (1 μg) was converted to cDNA by random hexamers using Verso c-DNA Kit (Thermo Fisher Scientific) following manufacturers’ protocol. The cDNA was then used in qPCR as template and RNA was quantified using SYBR Green chemistry. Gene expression was normalized to *sigA* (MSMEG_2758) and quantified by the ΔΔCt algorithm. Appropriate controls (without reverse transcriptase or template negative control and genomic DNA-positive) were incorporated in each experiment. Primers were designed using OligoAnalyzer™ Tool (IDT) and are listed in Table S1.

The RNAseq was outsourced from Medgenome Pvt. Ltd. RNA was isolated from three biological replicates of each strain. The isolated RNA was fragmented, cDNA library was prepared and RNAseq was performed. The raw reads were filtered using Trimmomatic for quality scores and adapters. Filtered reads were aligned to the *Mycobacterium smegmatis* genome using HISAT2 to quantify reads mapped to each transcript. Alignment percentages to the *Mycobacterium smegmatis* genome were in the range of 75-87.7 %. Total number of uniquely mapped reads were counted using feature counts. The uniquely mapped reads were used to analyse differential gene expression using DeSeq2. The ratio of normalized read counts for treated over control was taken as the fold change. Genes were first filtered based on the *p*-value (<= 0.05). The log2 (foldchange) values were found to be normally distributed. Genes showing change in expression within −1 ≤log2 (foldchange) ≥ 1 were considered.

### Growth curve assay

The recombinant cells grown to mid-log phase were used to inoculate fresh Middlebrook 7H9 medium with or without ATc at a starting O.D_600nm_ of 0.02 in a 96 well plate. The O.D_600nm_ was monitored in Tecan Pro plate reader at 37°C, every 2 hours for 48 hours. The O.D values were plotted against time to evaluate the fitness of strains.

### Electron microscopy

The knockdown strains (5 ml of 0.6 O.D cells) were harvested at 5000*g* for 10 min at 4 °C and washed twice in 0.1M phosphate buffered saline, pH-8 (PBS). The cells were then fixed using 2.5% glutaraldehyde for 2 hours in dark. The fixed cells were then centrifuged and washed with PBS to remove traces of glutaraldehyde. The washed cells were subsequently dehydrated by serially treating with alcohol concentrations: 10%, 30%, 40%, 50%, 70% and 90% respectively. The cells were finally suspended in 100% alcohol and air dried to remove excess water content from the sample. The sample was drop cast on a carbon taped aluminium stub and was imaged after platinum sputter coating. SEM was performed at ×10,000 on a FE-SEM (FEI-Quanta 200 at SAIF, IIT Bombay) instrument using an accelerating voltage of 20 kV.

### Cell permeabilisation assay

MSMEG_0311K.D and NTA strains were pre-depleted of targeted gene expression in presence of ATc for at least 15 hours. Subsequently, the cells were harvested and washed in PBS to remove any trace of media components. The cells were maintained at equal O.D of 0.6 and were treated with ethidium bromide at 2μg/ml or nile red at 20μM. The cells were incubated in 96-black well plate and absorbance at 600 nm and fluorescence (excitation: 530nm and emission: 590nm) were measured at 3-minute interval. The values were plotted against time.

### Disc assay

The knockdown strains were pre-depleted of target gene product in presence of ATc for at least 15 hours. Subsequently, the strains were harvested and washed in PBS (0.1M, pH8). The cells (200 μl at 0.D 1) were then suspended in warm 0.6% agarose and mixed well such that a homogenous suspension formed. The re-suspended cells were gently poured over MB agar plates containing hygromycin, kanamycin and ATc. The discs (ethambutol and isoniazid - 150μg, streptomycin, imipenem, and gentamycin-10μg, vancomycin, linezolid, and amikacin-30μg) impregnated with desired chemicals were placed gently on to the agar. The plates were incubated for 3-4 days until a visible lawn appeared and zone of inhibition was measured and tabulated.

### Antibiotic sensitivity Broth assay

All antibiotic compounds were dissolved in water. The growth-synchronised and pre-depleted knockdown strains were inoculated at O.D 0.02 in presence of ATc and drugs at applicable concentration. 200 μl culture was dispensed per well in a 96-well micro-titre plate in triplicates. O.D_600nm_ was evaluated using a Tecan Pro plate reader for 48 hours. A suitable time point was selected at which both strains were in the log phase for further analysis. Percent growth was calculated relative to the vehicle control for each strain for all curves, data represent the mean ± sem for technical triplicates. Data are representative of at least two independent experiments.

### Protein localisation assay

For protein localization studies, the cells expressing the GFP-fused protein were harvested at mid-log phase and washed twice in PBS (0.1M, pH-8). The washed cells were drop cast on a glass slide and imaged by confocal microscopy (Olympus Confocal microscope, FV030S, NIRRH)

### RADA pulse chase experiment

Saturated cultures of *M. smegmatis* strains bearing MSMEG_0311 GFP fused plasmid (pMV261-0311-GFP), MSMEG_0311 spacer (pSTHT-0311 K.D) or *mmpl spacer* (pSTHT-0250 K.D) were sub-cultured and grown to reach log phase to ensure all cells are in same phase of growth. Subsequently, cells were inoculated into fresh medium containing RADA dye at 20µM concentration at 0.02 O.D_600nm_. The cells were given a long pulse of 15 hours to ensure that all cells are uniformly stained with the dye, followed by addition of ATc for induction of CRISPRi. The cells were grown for 5 hours to allow depletion of the target genes. Cultures that did not receive ATc served as control for CRISPRi. Subsequently, the cells were washed thrice with media and allowed to grow in absence of RADA for 3 hours during the chase period. The cells were then washed with PBS and mounted on slides using Antifade (Invitrogen). The slides were visualised using TRITC filter (excitation: 560nm and emission: 630nm) using Olympus confocal microscope.

## Acknowledgement

Authors thank Snehalata Desai for the technical help in conduct of experiments. pMV261 was a gift from Dr. Amit Singh. pSTKiT was a kind gift from Dr. Vinay K. Nandicoori. pMN437 was a gift from Dr. Michael Niederweis (Addgene plasmid #32362). Authors thank SAIF, IITB facility for access of electron microscopy and NIRRH, Parel for access of confocal microscopy.

