## Supplementary material for "MSMEG_0311 is a conserved essential polar protein involved in mycobacterium cell wall metabolism"

### Supplementary information

**Table S1.** Sequence conservation between the homologues of MSMEG_0311 and neighbouring proteins

| Homologue in *M. tuberculosis* H37Rv | *M. bovis* | | *M. leprae* | | *M. smegmatis* | |
|---|---|---|---|---|---|---|
| | Identity | Similarity | Identity | Similarity | Identity | Similarity |
| *Rv0225c* | 100 | 100 | 85 | 91 | 76 | 85 |
| *Rv0226c* | 100 | 99 | 70 | 79 | 65 | 74 |
| *Rv0227c* | 99 | 99 | 76 | 85 | 65 | 78 |
| *Rv0228c* | 100 | 100 | 79 | 85 | 74 | 83 |

**Fig. S1** Sequence of dead-Cas12a codon optimised for Corynebacterium expression

```
    atgtcg atctaccaag agttcgtgaa taagtatagc ctgagcaaga ccttgcggtt
cgagttgatc ccgcagggca aaaccttgga aaacattaag gcgcgcgggc tgatcctgga
tgatgagaag cgggccaagg actataaaaa ggccaagcag attatcgata aatatcatca
gttcttcatc gaggaaatcc tgtcgagcgt gtgcatcagc gaagacttgt tgcagaacta
tagcgacgtg tacttcaaat tgaaaaagtc ggacgacgac aatctgcaaa aagacttcaa
gtcggccaaa gacaccatca agaaacagat ttcggagtac atcaaggact cggaaaagtt
caaaaacctg ttcaaccaga acttgatcga cgcgaaaaag ggccaggaaa gcgatttgat
cttgtggctg aagcagtcga aggataatgg cattgagttg ttcaaggcca actcggacat
tacggacatt gacgaggccc tggaaatcat taaatcgttt aagggctgga ccacgtactt
caagggcttc catgagaacc ggaaaaacgt gtattcgtcg aacgacattc ccacgtcgat
catctaccgg atcgtcgatg acaacttgcc caagttcttg gaaaataaag cgaaatacga
gagcttgaag gacaaagcgc cggaggccat taactatgag cagatcaaga aggacctggc
cgaagagctg acctttgaca ttgactacaa gaccagcgag gtcaatcagc gggtcttctc
gttggatgag gtctttgaaa tcgccaattt caacaactat ttgaaccaga gcggcatcac
caaatttaat accattatcg gcgggaagtt cgtcaacggc gagaacacga agcggaaggg
gatcaacgaa tacatcaatc tgtactcgca gcagatcaac gacaagaccc tgaaaaagta
caagatgagc gtgctgttca agcaaattct gagcgacacc gagagcaagt cgttcgtcat
tgacaagctg gaggatgaca gcgatgtggt caccacgatg cagtcgtttt acgagcaaat
cgcggccttc aagaccgtgg aggaaaagag catcaaggaa accctgagcc tgctgttcga
tgatctgaag gcgcagaagt tggatctgtc gaaaatctac ttcaagaacg ataagtcgct
gaccgacctg agccaacagg tcttcgatga ttacagcgtc atcgggaccg ccgtcctgga
gtacatcacc cagcagattg cgcccaaaaa cctggacaat ccctcgaaaa aagagcagga
gctgattgcc aagaagaccg agaaggcgaa gtacttgtcg ctggaaacca tcaaactggc
gctggaggag ttcaacaagc accgcgacat tgataaacaa tgccggttcg aggagattct
ggccaatttt gccgccattc ccatgatctt cgatgagatc gcgcagaata aagacaacct
ggcgcaaatc tcgatcaaat atcagaatca gggcaaaaaa gacctgttac aagcctcggc
cgaagacgac gtgaaagcca tcaaggacct gttggatcag acgaacaact tgttgcacaa
gctgaaaatt ttccacatca gccagagcga ggacaaggcc aacatcttgg acaaagacga
```

gcacttctac ttggtgttcg aagagtgcta cttcgaattg gccaacatcg tccccctgta
caataagatt cgcaattata ttacgcagaa gccctatagc gacgagaaat tcaagttgaa
ttttgaaaat tcgaccctgg cgaatggctg ggacaagaac aaagagcccg acaacaccgc
catcttgttc atcaaggatg ataaatatta tctgggggtg atgaacaaaa agaataacaa
gatctttgat gacaaggcca tcaaggagaa taagggcgag ggtacaaga aaatcgtcta
caaactgttg cccggcgcca acaagatgtt gccgaaggtc ttcttcagcg cgaagagcat
taaattctat aaccccagcg aggacatcct gcgcatccgg aatcattcga cccacacgaa
gaatggcagc ccgcaaaagg gctacgagaa gttcgagttt aacatcgagg attgtcggaa
attcatcgac ttctataaac agtcgatcag caagcacccc gagtggaagg atttcggctt
ccgcttcagc gatacccaac gctacaacag catcgacgag ttttatcggg aagtggaaaa
ccaggggtac aagttgacct tcgagaacat ttcggagtcg tacattgact cggtcgtcaa
ccaagggaaa ctgtacctgt tccaaatcta caataaggac ttcagcgcgt actcgaaagg
gcggcccaac ttgcacaccc tgtattggaa ggccttgttt gacgagcgga acttacaaga
cgtcgtctat aaactgaacg gggaggcgga gctgttttac cgcaagcagt cgatcccgaa
aaagattacc cacccggcga aggaggcgat cgccaacaaa aacaaggaca acccgaagaa
agagtcggtc ttcgaatacg atctgatcaa agacaagcgc tttaccgagg acaagttctt
tttccattgt ccgatcacca tcaattttaa gagcagcggc gcgaacaagt tcaacgacga
gatcaacctg ctgttgaaag aaaaggccaa tgatgtgcac atcctgtcga tcgcccgggg
ggaacggcac ctggcctact acaccctggt ggacggcaag ggcaacatca tcaagcagga
taccttcaat atcattggga acgaccggat gaaaacgaac taccacgaca agctggcggc
catcgaaaag gatcgcgaca gcgcccgcaa ggactggaag aagattaaca atatcaagga
gatgaaggag gggtacttga gccaggtcgt gcacgagatc gccaagctgg tcattgagta
caacgcgatt gtggtctttg aagacctgaa ctttggcttc aaacgggggc ggttcaaggt
cgagaagcag gtgtatcaga agctggaaaa aatgttgatc gagaagctga actatctggt
gttcaaagac aacgagtttg acaaaacggg gggcgtcctg cgggcctacc agttgaccgc
cccttcgaa accttcaaga aaatgggcaa gcagacgggc atcatctact acgtcccggc
gggctttacg agcaagatct gtccggtgac cggcttcgtc aaccagctgt atccgaagta
tgagagcgtg tcgaaaagcc aggagttttt ctcgaagttc gataaaatct gttataacct
ggataagggg tacttcgaat tttcgttcga ttacaaaaac ttcggggaca aggcggccaa
ggggaaatgg acgatcgcca gcttcggctc gcgcctgatt aactttcgca actcggacaa
gaaccataac tgggacaccc gggaagtgta cccgaccaaa gaattggaaa aactgttgaa
ggactatagc atcgagtacg ggcacgggga gtgcatcaaa gccgccatct gcggggaaag
cgacaagaag ttttttgcga agctgaccag cgtgctgaat acgatcttac aaatgcgtaa
ttctaagacc ggcaccgagc tggactacct gatctcgccg gtggcggatg tcaacggcaa
cttcttcgac agccgccagg ccccgaagaa catgccgcag gacgcggatg cgaatggcgc
gtaccacatc gggctgaaag ggctgatgct gctggggcgg atcaagaaca accaggaggg
gaagaagctg aatctggtga ttaagaacga ggaatatttc gagttcgtcc aaaatcgcaa
caattga

**The codon mutagenized (D917A) to convert Cas12a into dead-Cas12a is highlighted.**

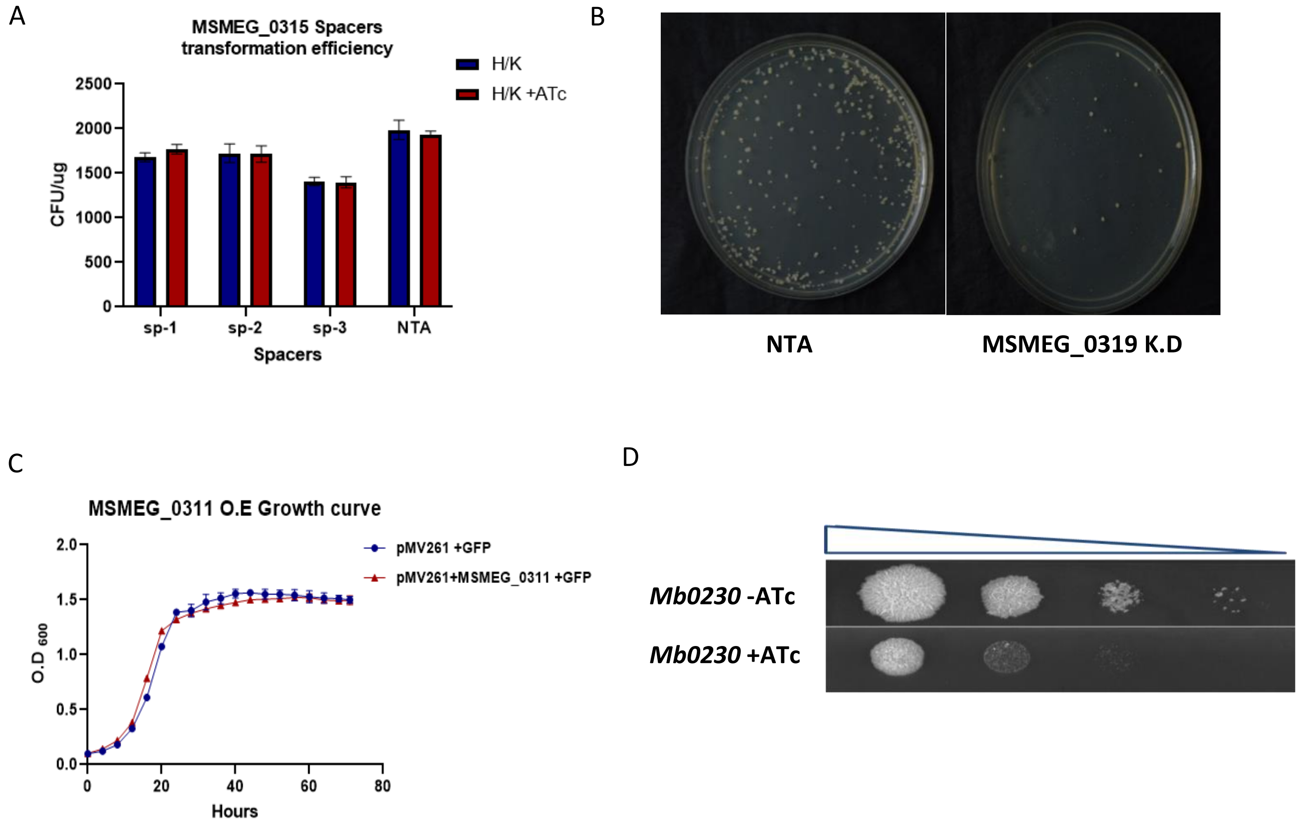


**Fig. S2 A)** Transformation efficiency of plasmids expressing gRNA containing spacers targeting SMEG_0315 **B)** Growth curve of *M. smegmatis* over-expressing (O.E) MSMEG_0311 **C)** Colony morphology of MSMEG_0319 KD strain and NTA **D)** MSMEG_0311 homologue Mb0230 is essential in *M.bovis* BCG

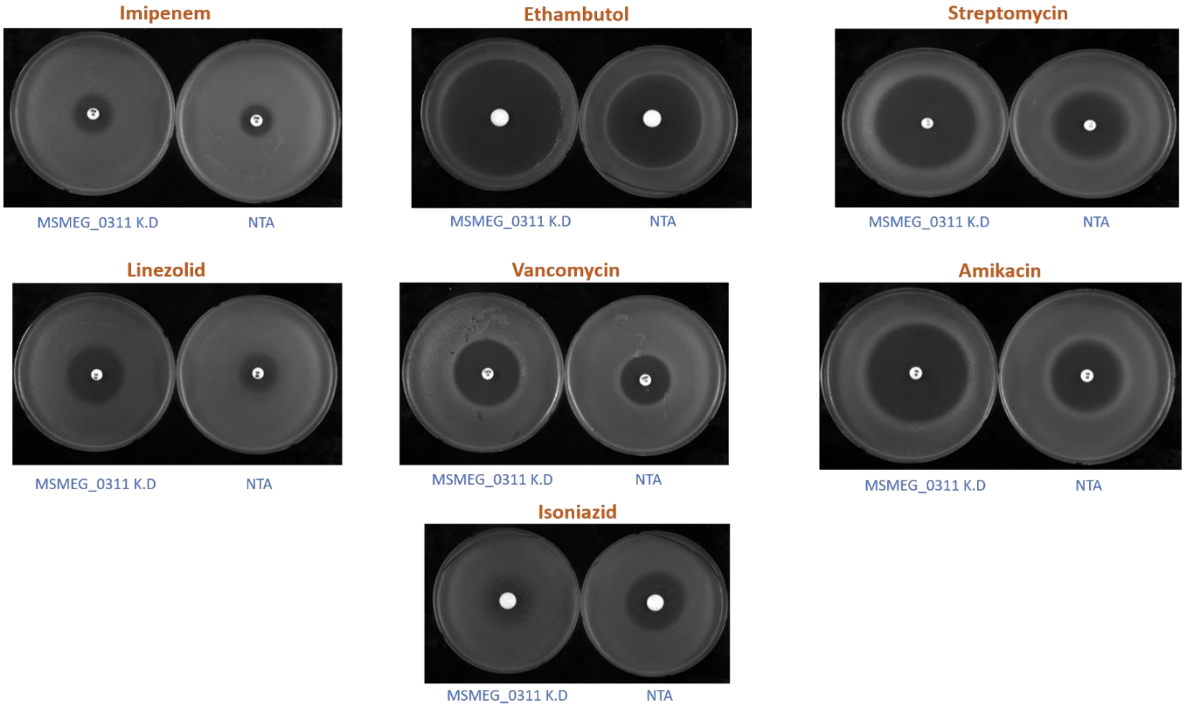


**Fig. S3** Antibiotic susceptibility assay of MSMEG_0311 K.D and NTA

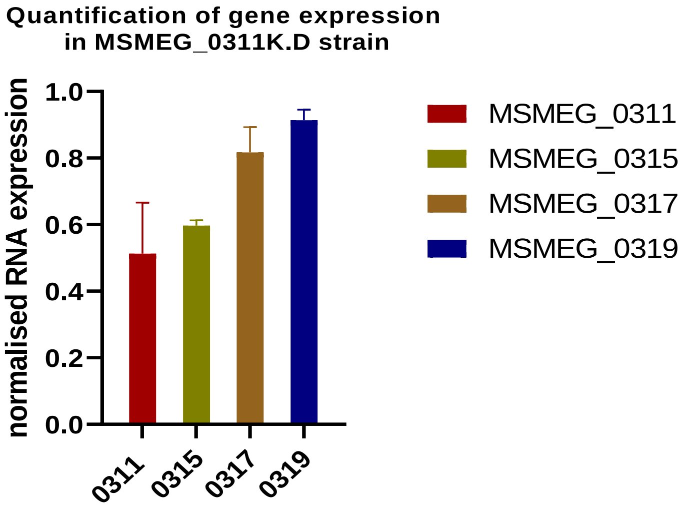


**Fig. S4** Quantification of expression of genes with qPCR

**Table S2** List of plasmids used in the study

| S.No | Plasmid name | Description | Reference |
|---|---|---|---|
| 1. | pSTKiT | Mycobacterium integrative shuttle vector | Parikh et al., 2013 |
| 2. | pSTKiT-Cas12 | PSTKiT backbone with optimised tetO promoter and codon optimised dead *cas12* cloned at KpnI and HindIII site | This study |
| 3. | pSTHT | Mycobacterium replicative shuttle vector with $Hyg^R$ | This study |
| 4. | pSTHT _0310 K.D | pSTHT backbone with spacer cloned against MSMEG_0310 at BsaI site | This study |
| 5. | pSTHT _0311 K.D | pSTHT backbone with spacer cloned against MSMEG_0311 at BsaI site | This study |
| 6. | pSTHT_0315 K.D | pSTHT backbone with spacer cloned against MSMEG_0315 at BsaI site | This study |
| 7. | pSTHT _0317 K.D | pSTHT backbone with spacer cloned against MSMEG_0317 at BsaI site | This study |
| 8. | pSTHT_0319 K.D | pSTHT backbone with spacer cloned against MSMEG_0319 at BsaI site | This study |
| 9. | pSTHT _0250 K.D (*mmpl*) | pSTHT backbone with spacer cloned against MSMEG_0250 at BsaI site | This study |
| 10. | pMV261 | Mycobacterium replicative shuttle vector with $Kan^R$ | Stover et al., 1991 |
| 11. | pMV261-0311 | pMV261 with MSMEG_0311 cloned at BamHI and EcoRI site | This study |
| 12. | pMV261 –0311-GFP | pMV261 -0311 with GFP cloned at EcoRI and HindIII site | This study |

**Table S3** List of oligonucleotides used in the study

| S.No | Oligo name | Sequence 5’-3’ | Description |
|---|---|---|---|
| 1. | Msm_310F | TAGATGCCGGCCGGGCGACGCTCCG | Spacer used to target MSMEG_0310 in CRISPRi |
| 2. | Msm_310R | AGACCGGAGCGTCGCCCGGCCGGCA | Spacer used to target MSMEG_0310 in CRISPRi |
| 3. | Msm_311 F | TAGATGGCGCGCACCGAGGTGGTCG | Spacer used to target MSMEG_0311 in CRISPRi |
| 4. | Msm_311 R | AGACCGACCACCTCGGTGCGCGCCA | Spacer used to target MSMEG_0311 in CRISPRi |
| 5. | Msm_315 F1 | TAGATACGGCCGCGACGCTGGCGAT | Spacer used to target MSMEG_0315 in CRISPRi |

| | | | |
|---|---|---|---|
| 6. | Msm_315 R1 | AGACATCGCCAGCGTCGCGGCCGTA | Spacer used to target MSMEG_0315 in CRISPRi |
| 7. | Msm_315 F2 | TAGATGATGCTGGTCTGGCCGGCCG | Spacer used to target MSMEG_0315 in CRISPRi |
| 8. | Msm_315 R2 | AGACCGGCCGGCCAGACCAGCATCA | Spacer used to target MSMEG_0315 in CRISPRi |
| 9. | Msm_315 F3 | TAGATATGCCGCAGAGCGGTATGGGCAT | Spacer used to target MSMEG_0315 in CRISPRi |
| 10. | Msm_315 R3 | AGACATGCCCATACCGCTCTGCGGCATA | Spacer used to target MSMEG_0315 in CRISPRi |
| 11. | Msm_317 F | TAGATAACCGCGCTGTGGCGCTGCG | Spacer used to target MSMEG_0317 in CRISPRi |
| 12. | Msm_317 R | AGACCGCAGCGCCACAGCGCGGTTA | Spacer used to target MSMEG_0317 in CRISPRi |
| 13. | Msm_319F | TAGATCTGCCCGCGGTCGAAGGCATG | Spacer used to target MSMEG_0319 in CRISPRi |
| 14. | Msm_319 R | AGACCATGCCTTCGACCGCGGGCAGA | Spacer used to target MSMEG_0319 in CRISPRi |
| 15. | Msm_0250 F | TAGAT GCCTGGTGGGGTCGGACCGTG | Spacer used to target *mmpl* in CRISPRi |
| 16. | Msm_0250 R | AGACCACGGTCCGACCCCACCAGGCA | Spacer used to target *mmpl* in CRISPRi |
| 17. | Nta 1F | TAGATACCAAGGACACATTCGAGCTCT | Scrambled nn targeting spacer used as control |
| 18. | Nta 1R | AGACAGAGCTCGAATGTGTCCTTGGTA | Scrambled nn targeting spacer used as control |
| 19. | Msmeg 6766 RTF | GCTGTCCGTGATCTCGACC | qRT PCR Primer used to validate RNASeq data for MSMEG_6766 expression |
| 20. | Msmeg 6766 RTR | CATCTGCACGGTCTCACCC | qRT PCR Primer used to validate RNASeq data for MSMEG_6766 expression |
| 21. | Msmeg 2343 RTF | GAGGCCAAACCCGCGATC | qRT PCR Primer used to validate RNASeq data for MSMEG_2343 expression |
| 22. | Msmeg 2343 RTR | GAGTTCGTCGAGGCGCAC | qRT PCR Primer used to validate RNASeq data for MSMEG_2343 expression |
| 23. | Msmeg 4391 RTF | CCATCACGTACGGCTCACAAC | qRT PCR Primer used to validate RNASeq data for MSMEG_4391expression |
| 24. | Msmeg 4391 RTR | CTCCTCGATGTGGGCGAAGT | qRT PCR Primer used to validate RNASeq data for MSMEG_4391 expression |
| 25. | Msmeg 6606 RTF | GGTGACGGTGCTGATCATCG | qRT PCR Primer used to validate RNASeq data |

| | | | |
|---|---|---|---|
| | | | for MSMEG_6606 expression |
| 26. | Msmeg 6606 RTR | GTGTCGGCGATTCCCAGC | qRT PCR Primer used to validate RNASeq data for MSMEG_6606 expression |
| 27. | Msmeg_6767 RTF | CGCGTGAGGTACCGGAG | qRT PCR Primer used to validate RNASeq data for MSMEG_6767 expression |
| 28. | Msmeg_6767 RTR | CACGTCGGTACCTGCGAG | qRT PCR Primer used to validate RNASeq data for MSMEG_6767 expression |
| 29. | Msmeg 6583 RTF | CGCTGTCACGTCGTGTCG | qRT PCR Primer used to validate RNASeq data for MSMEG_6583 expression |
| 30. | Msmeg 6583 RTR | AAC GCC GCG GTG TTG ATG | qRT PCR Primer used to validate RNASeq data for MSMEG_6583 expression |
| 31. | Msmeg 0695 RTF | GATCCGCAGATCCGCGTC | qRT PCR Primer used to validate RNASeq data for MSMEG_0695 expression |
| 32. | Msmeg 0695 RTR | GCCACCTTTGAGCAGCGG | qRT PCR Primer used to validate RNASeq data for MSMEG_0695 expression |
| 33. | Msmeg 2016 RTF | GCCTCGCTCAAGTCGGTG | qRT PCR Primer used to validate RNASeq data for MSMEG_2016 expression |
| 34. | Msmeg 2016 RTR | CTTCTCGGTCGCCGCTC | qRT PCR Primer used to validate RNASeq data for MSMEG_2016 expression |
| 35. | pMV seqR | TGATGCCTGGCAGTCGATC | Primer used to sequence 311GFP fusion construct |
| 36. | 311 BamH1 F | TATGGATCCATGTCTGCCCGGCCCGAGTC TGCTCCG | Primer used to amplify and clone 0311 for GFP fusion construct |
| 37. | 311 linker R | TTAGAATTCGCCAGAACCAGCAGCGGAG CCAGCCGAGACCAGGCCGCTGAGGTATC | Primer used to amplify and clone 0311 for GFP fusion construct |
| 38. | Gfp F | TTATGAATTCATGTCGAAGGGCGAGGAG CT | Primer used to amplify and clone GFP for GFP fusion construct |
| 39. | Gfp flag R | TATAAGCTTCTACTTGTCGTCGTCGTCCTT GTAGTCCTTGTACAGCTCGTCCAT | Primer used to amplify and clone GFP for GFP fusion construct |
| 40. | 0317 RTF | GTCCACGCTCATCGATCC | qRT PCR Primer used to quantitate MSMEG_0317 gene expression |
| 41. | 0317 RTF | GGCAGCTTCTCGGTCTTG | qRT PCR Primer used to quantitate MSMEG_0317 gene expression |

| | | | |
|---|---|---|---|
| 42. | 311 genotyp RTF | GTCACGTCGACGAGGACACC | qRT PCR Primer used to quantitate MSMEG_0311 gene expression |
| 43. | 311 genotyp RTR | CGGTAGCCGATGGTGGGAAC | qRT PCR Primer used to quantitate MSMEG_0311 gene expression |
| 44. | 315 genotyp RTF | GTGCTGCCGGATCTGACGTG | qRT PCR Primer used to quantitate MSMEG_0315 gene expression |
| 45. | 315 genotyp RTR | AGAACTCGCGGATGCTGTCG | qRT PCR Primer used to quantitate MSMEG_0315 gene expression |
| 46. | 319 genotyp RTF | GGGCTGTTCGTGTGGCATCT | qRT PCR Primer used to quantitate MSMEG_0319 gene expression |
| 47. | 319 genotyp RTR | GAAGCCGAACACCAGCGTGA | qRT PCR Primer used to quantitate MSMEG_0319 gene expression |
| 48. | Mys genotyp RTF | CCGAAGAGCTCGCCAAGGAG | qRT PCR Primer used to normalise differential gene expression |
| 49. | Mys genotyp RTR | GTCACCGAGCTGGCTGTCAC | qRT PCR Primer used to normalise differential gene expression |
